# Immature olfactory sensory neurons provide complementary input in the healthy olfactory system

**DOI:** 10.1101/2025.11.05.686656

**Authors:** Jordan D. Gregory, Ryan S. Herzog, Kendall A. Curtis, Michael I. Marar, Claire E.J. Cheetham

## Abstract

Adult neurogenesis of olfactory sensory neurons (OSNs) provides the unique opportunity to understand how new neurons functionally integrate into existing circuitry and contribute to behaviors. Previous studies have shown that immature OSNs express odorant receptors, form dendritic knobs with short cilia, and project their axons into the olfactory bulb (OB) to form functional synapses. Furthermore, a previous study found that immature OSNs respond selectively to odorants and exhibit graded responses in a higher odorant concentration range than mature OSNs. Finally, this study also showed that, in mice that lack mature OSNs, sensory input from immature OSNs can mediate odor detection and discrimination. What remains unknown is how these immature OSNs contribute to odor-guided behavior in the healthy, intact olfactory system. Here we show that chemogenetically silencing immature OSNs impairs odor detection ability in a buried food assay. Furthermore, immature OSN silencing reduces the amplitude of odor-evoked dendritic calcium responses in OB neurons *in vivo*. Together, these findings suggest that immature OSNs provide distinct odor input that complements mature OSN input to contribute to odor-guided behaviors in the healthy, intact olfactory system.

## Introduction

Most mammalian neurons are generated embryonically or early in postnatal development and cannot be replaced. In contrast, newborn olfactory sensory neurons (OSNs) are generated from basal stem/progenitor cells in the olfactory epithelium (OE) throughout life in all terrestrial mammals, including humans^1,2^. Horizontal basal cells are multipotent stem cells that are normally quiescent but undergo mitosis in response to OE injury^3–6^. Globose basal cells are progenitor cells that undergo constitutive mitosis in the healthy OE to generate both sustentacular cells (OE support cells) and OSNs and can themselves be generated from HBCs during OE regeneration after injury^3,7,8^. Terminal cell division of GBCs produces neuronally committed intermediate cells that undergo a developmental program to produce mature OSNs^7–10^. One to two days after this terminal cell division, nascent OSNs transition to immature OSNs marked by the expression of both GAP43 and Gγ8^9,11–15^. Seven to eight days after terminal division, OSNs downregulate their expression of GAP43 and Gγ8 and upregulate expression of olfactory marker protein (OMP)^1,2,14,16–18^.

In mice, mature OSNs typically express one of ∼1,100 odorant receptor (OR) genes^19,20^, with mature OSN axons expressing the same OR converging into the same glomeruli in the olfactory bulb (OB)^21–23^. Here, they form synaptic connections with OB neurons, generating a highly organized odor input map. As existing OSNs undergo apoptosis, newborn OSNs are generated concurrently. This constant turnover creates a highly plastic environment in which newborn OSNs must integrate into pre-existing circuitry to contribute to olfaction. Hence, it is important to understand how ongoing OSN neurogenesis contributes to odor processing, plasticity, and odor-guided behavior.

It has been assumed that mature OSNs provide the sole source of odor input to the OB. However, mounting evidence suggests that immature OSNs also contribute to olfaction. OSNs begin to express ORs four days after terminal cell division^14^, well before the onset of OMP expression. Furthermore, immature OSNs strongly upregulate neurite outgrowth genes^10^ and begin to both project axons into glomeruli in the OB and form dendritic knobs with short cilia^24–28^. Moreover, immature OSNs can detect odorants and form synaptic connections with OB projection neurons in mice^25,27^. Importantly, odor selectivity is similar for immature and mature OSNs^27^. However, compared to mature OSNs, immature OSNs have higher thresholds for odorant detection but continue to encode concentration information at high odorant concentrations at which mature OSN responses are saturated^27^. Furthermore, sensory input from immature OSNs is sufficient to mediate odor detection and discrimination at early recovery time points in mice, just 5–7 days after they have undergone methimazole-mediated OSN ablation^27,29,30^. Together, these data suggest that immature OSNs provide behaviorally relevant information to the olfactory system.

What remains unknown is what contribution, if any, immature OSNs make in the healthy, intact olfactory system. We hypothesize that immature and mature OSNs provide complementary, but distinct, odor input to OB neurons. Here, we used selective DREADD (designed receptors exclusively activated by designer drugs)-based chemogenetic silencing^31–33^ of immature OSNs to address two key questions. First, do immature OSNs contribute to odor-guided behaviors in healthy mice? Second, do immature OSNs provide odor input to the OB that cannot be supplied by mature OSNs? We found that silencing immature OSNs impairs odor detection ability. Furthermore, acutely silencing immature OSNs reduces the amplitude of odorant-evoked responses in OB neurons *in vivo*. Together, our data suggest that immature and mature OSNs provide complementary odor input to the OB that contributes to distinct aspects of odor-guided behavior.

## Results

### Selective expression of hM4Di silences ∼60% of immature OSNs

To determine the contribution of immature OSNs to olfaction in the healthy, intact olfactory system, we employed chemogenetic silencing using hM4Di, an inhibitory DREADD that is active in the presence of a ligand such as clozapine-N-oxide (CNO). To achieve selective silencing of immature OSNs, we bred Gγ8-tTA;tetO-hM4Di (referred to hereafter as Gγ8-hM4Di) mice. In this line, expression of the tetracycline transactivator under control of the Gγ8 promoter, which is expressed only in immature OSNs, drives expression of hM4Di from a tetO (tTA-dependent) promoter (Fig. 1A). Hence, in the presence of CNO, hM4Di-expressing immature OSNs will be selectively silenced. tetO-hM4Di mice, which do not express hM4Di due to the lack of a tTA driver, served as controls. We validated the specificity of hM4Di expression in immature OSNs in Gγ8-hM4Di mice via co-immunostaining for GAP43 to mark immature OSNs and the hemagglutinin (HA) tag attached to the hM4Di receptor (Fig. 1B). tetO-hM4Di mice were included as negative controls throughout this study and did not express the hM4Di receptor marked by HA staining (Fig. 1C). Approximately 60% of immature OSNs expressed the hM4Di receptor in Gγ8-hM4Di mice, whereas tetO-hM4Di controls, which lack tTA expression, did not express hM4Di (Fig. 1D).

**Figure 1.**
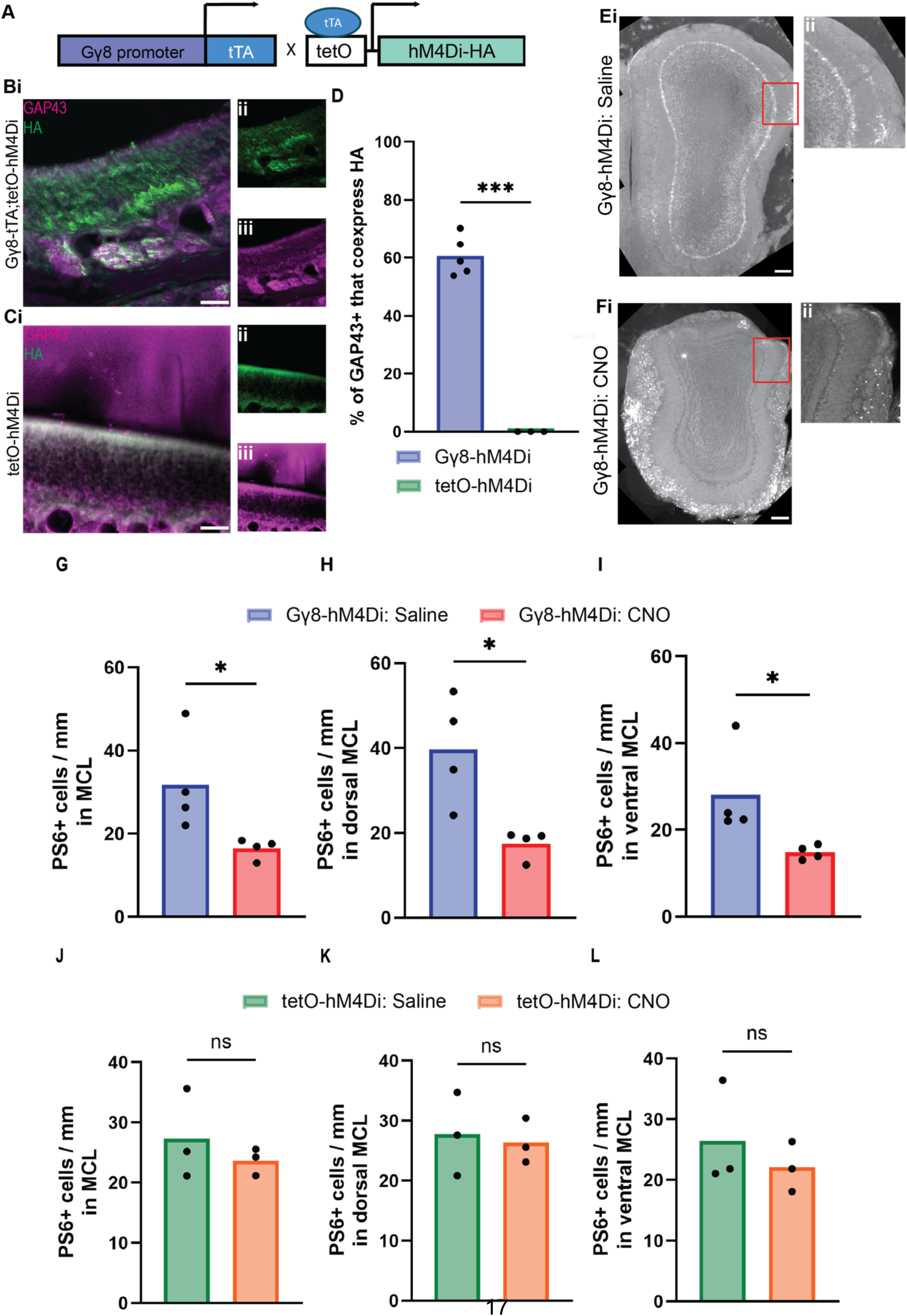
Validation of transgenic approach to silence immature OSNs. **A.** Breeding strategy to generate Gγ8-tTA;tetO-hM4Di [Gγ8-hM4Di] experimental and tetO-hM4Di control mice. **B,C. i.** 60x magnification maximum intensity projection confocal images of OE stained for HA (green) and GAP43 (magenta) in a Gγ8-hM4Di mouse (**B**) and a tetO-hM4Di mouse (**C**). Scale bars: 100 µm. **ii.** Single channel images showing HA staining. **iii.** Single channel images showing GAP43 staining. **D.** Percentage of GAP43-positive OSNs that co-express the hM4Di receptor marked with the HA tag (Gγ8-hM4Di n = 5; tetO-hM4Di n = 3; Welch’s t-test: *t*(4) = 19.9, *P* < 0.001). **E,F. i.** Example images of OBs from a saline-treated Gγ8-hM4Di mouse (**E**) and a CNO-treated Gγ8-hM4Di mouse (**F**) stained for PS6. Red box outlines area of zoomed panels (right of images) used for counting PS6+ cells shown in **ii**. Scale bars: 100 µm. Note that bright fluorescent puncta in GL of panel F are due to lipofuscin autofluorescence. **G.** Linear density of PS6+ cells in the MCL is decreased by silencing immature OSN input (unpaired t-test: *t*(6) = 2.54, *P* = 0.044, n = 4 mice per group). **H,I.** Reduced PS6+ MC density when immature OSNs are silenced during odor exposure in dorsal (**H**) (unpaired t-test: *t*(6) = 3.35, *P* = 0.015) and ventral (**I**) (unpaired t-test: *t*(6) = 2.46, *P* = 0.049) MCL. **J.** Linear density of PS6+ cells in the MCL is not affected by CNO administration in tetO-hM4Di control mice (unpaired t-test: *t*(3) = 0.81, *P* = 0.46, n = 3 mice per group). **K, L.** No effect of CNO administration during odor exposure on linear density of PS6+ MCs in dorsal (**K**) (unpaired t-test: *t*(3) = 0.30, *P* = 0.78) or ventral (**L**) (unpaired t-test: *t*(3) = 0.79, *P* = 0.47) MCL in tetO-hM4Di control mice (n = 3 mice per group). Symbols: values for individual mice.

### Silencing immature OSNs reduces mitral cell activity in response to odors throughout the OB

We first sought to validate our model to acutely silence immature OSNs and determine the effects of immature OSN silencing on odor-evoked mitral cell (MC) activity. We stained the OB for the immediate early gene phospho-S6 (PS6) in saline- and CNO-treated Gγ8-hM4Di mice after a 1-hour odor exposure session (Fig. 1E,F). We then calculated the linear density of PS6+ cells in the mitral cell layer (MCL) for each mouse. This showed that silencing immature OSNs resulted in a significantly lower density of PS6+ MCs, indicating that MC activity was reduced as a result of immature OSN silencing (Fig. 1G; saline-treated Gγ8-hM4Di n = 4, CNO-treated Gγ8-hM4Di n = 4). It has been reported both that proliferation occurs at a higher rate in the ventral OE than the dorsal OE^17^, and that the ventral OE contains a larger proportion of immature OSNs than the dorsal OE^34^. It is also known that OE-OB projections are topographic, with dorsal zone OSNs projecting to the dorsal OB, while ventral zone OSNs project to the ventral OB^35,36^. Hence, it was possible that MC activity might be differentially affected by immature OSN silencing in the dorsal vs. ventral OB. Subdividing the MCL showed that immature OSN silencing had a significant effect on MC activity in both dorsal and ventral regions (Fig. 1H,I). The reduction in MC PS6 staining was not present in CNO-treated tetO-hM4Di control mice (Fig. 1J–L; saline-treated tetO-hM4Di n = 3, CNO-treated tetO-hM4Di n = 3), thus validating our strategy to acutely and selectively silence immature OSN input throughout the OB using CNO treatment in Gγ8-hM4Di mice.

### Immature OSN silencing impairs odor detection behavior

Odor detection ability was tested using a buried food assay wherein mice are given 10 minutes to complete the task (Fig. 2A). In Gγ8-hM4Di mice, all saline-injected mice successfully uncovered the buried food within the 10-minute period (n = 7 mice), whereas two out of eight CNO-injected Gγ8-hM4Di mice failed to uncover the buried food (Fig. 2B). In tetO-hM4Di control mice, three out of five saline-injected and all five CNO-injected mice uncovered the buried food (Fig. 2B). The time taken to uncover the buried food was significantly longer for CNO-treated than for saline-treated Gγ8-hM4Di mice (Fig. 2B). There were no other significant differences in time to uncover the buried food between experimental groups, including no effect of CNO treatment in tetO-hM4Di control mice (Fig. 2B). There was also no difference in the time spent digging during the five-minute acclimation period between the four experimental groups (Fig. 2C), indicating that CNO treatment did not alter motor function in Gγ8-hM4Di mice. We also observed that all mice across experimental groups ate the food after completion of the assay, indicating similar motivation to perform the task. Together, these data show that odor detection ability is impaired when immature OSNs are silenced.

**Figure 2.**
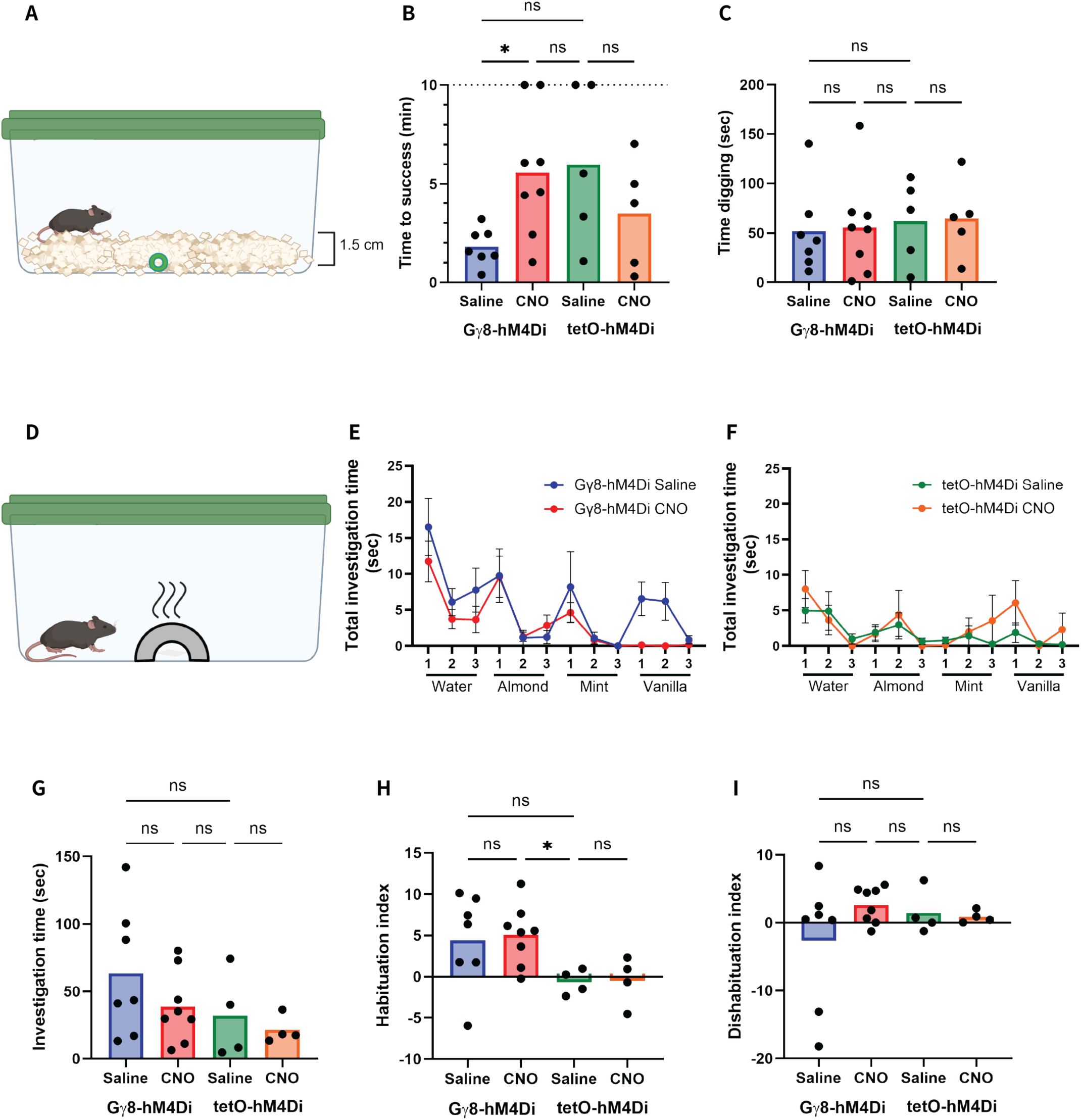
Immature OSNs contribute to odor detection ability in the intact olfactory system. **A.** Schematic of the buried food assay to test odor detection and source localization abilities. **B.** Acute silencing of immature OSN input via CNO injection in Gγ8-hM4Di mice results in increased time to success in buried food task (Welch’s one-way ANOVA: W(3, 8.63) = 4.44, *P* = 0.037. Dunnett’s T3 multiple comparisons tests: Gγ8-hM4Di saline vs. Gγ8-hM4Di CNO mice: t(8.31) = 3.16, *P* = 0.048; Gγ8-hM4Di saline vs. tetO-hM4Di saline mice: t(4.31) = 2.30, *P* = 0.24; Gγ8-hM4Di CNO vs. tetO-hM4Di saline treated mice: t(7.22) = 0.19, *P* = 1.00; tetO-hM4Di saline vs. tetO-hM4Di CNO mice: t(7.18) = 1.16, *P* = 0.69, saline-treated Gγ8-hM4Di mice n = 7, CNO-treated Gγ8-hM4Di mice n = 8, saline-treated tetO-hM4Di mice n = 5, CNO-treated tetO-hM4Di mice *n* = 5). Symbols: individual mice.

### Immature OSN silencing may not affect odor discrimination in a habituation-dishabituation task

To test whether odor discrimination ability is impacted by immature OSN silencing, we performed an odor habituation-dishabituation task, in which mice were presented with three consecutive trials each of four different odors (Fig. 2D-F). Mice that can discriminate the odors will both habituate to a familiar odor (sniff less to repeated presentations of the same odor) and dishabituate to a novel odor (sniff more when a novel odor is presented). We again tested saline- and CNO-treated Gγ8-hM4Di and tetO-hM4Di mice. We found no significant differences in investigation time between experimental groups (Fig. 2G).

For habituation index, we found no effect of CNO treatment in either Gγ8-hM4Di or tetO-hM4Di mice, although there was a significant difference between CNO-treated Gγ8-hM4Di and saline-treated tetO-hM4Di mice (Fig. 2H). For dishabituation index, there were no significant differences between experimental groups (Fig. 2I).

The lack of significant differences between saline- and CNO-treated Gγ8-hM4Di mice may suggest that immature OSN silencing does not affect odor discrimination in this assay. However, we also noted that some experimental groups did not exhibit strong evidence of habituation or dishabituation (Fig. 2E–I). To quantify this observation, we compared the habituation or dishabituation index for mice in each group to zero to determine whether they showed significant habituation or dishabituation, respectively. Significant habituation occurred only in saline- and CNO-treated Gγ8-hM4Di mice, and significant dishabituation occurred only in CNO-treated Gγ8-hM4Di mice (Fig. S1). Therefore, mice in some experimental groups did perform as expected in the odor habituation-dishabituation assay, and we cannot draw a strong overall conclusion about the effect of immature OSN silencing on odor discrimination.

### Immature OSN silencing reduces the amplitude of odor-evoked responses in OB neuron dendrites in the dorsal OB glomerular layer

To investigate the impact of immature OSN silencing on odor-evoked responses in OB neurons, we implanted acute cranial windows over the right OB of 3-5-week-old Gγ8-hM4Di and tetO-hM4Di mice that had received a neonatal injection of AAV5-syn-GCaMP6s directly into the right OB (Fig. 3A,B). This allowed us to record odor-evoked responses in the glomerular layer (GL) of the olfactory bulb *in vivo* before and after CNO injection. Neonatal AAV5-syn-GCaMP6s injection resulted in GCaMP6s expression in neuronal types throughout the OB, but most prominently in the MCL, external plexiform layer (EPL), and GL (Fig. 3C,D). Analysis of GCaMP6s-expressing cells in these OB layers revealed that a majority of the cells were found in the MCL (Fig. 3E) with apical dendrites of putative MCs and tufted cells (TCs) expressing GCaMP6s also projecting into the GL. Thus, MC and TC dendrites are likely to be a major contributor to the GL odor-evoked calcium responses that we report using AAV5-syn-GCaMP6s.

**Figure 3.**
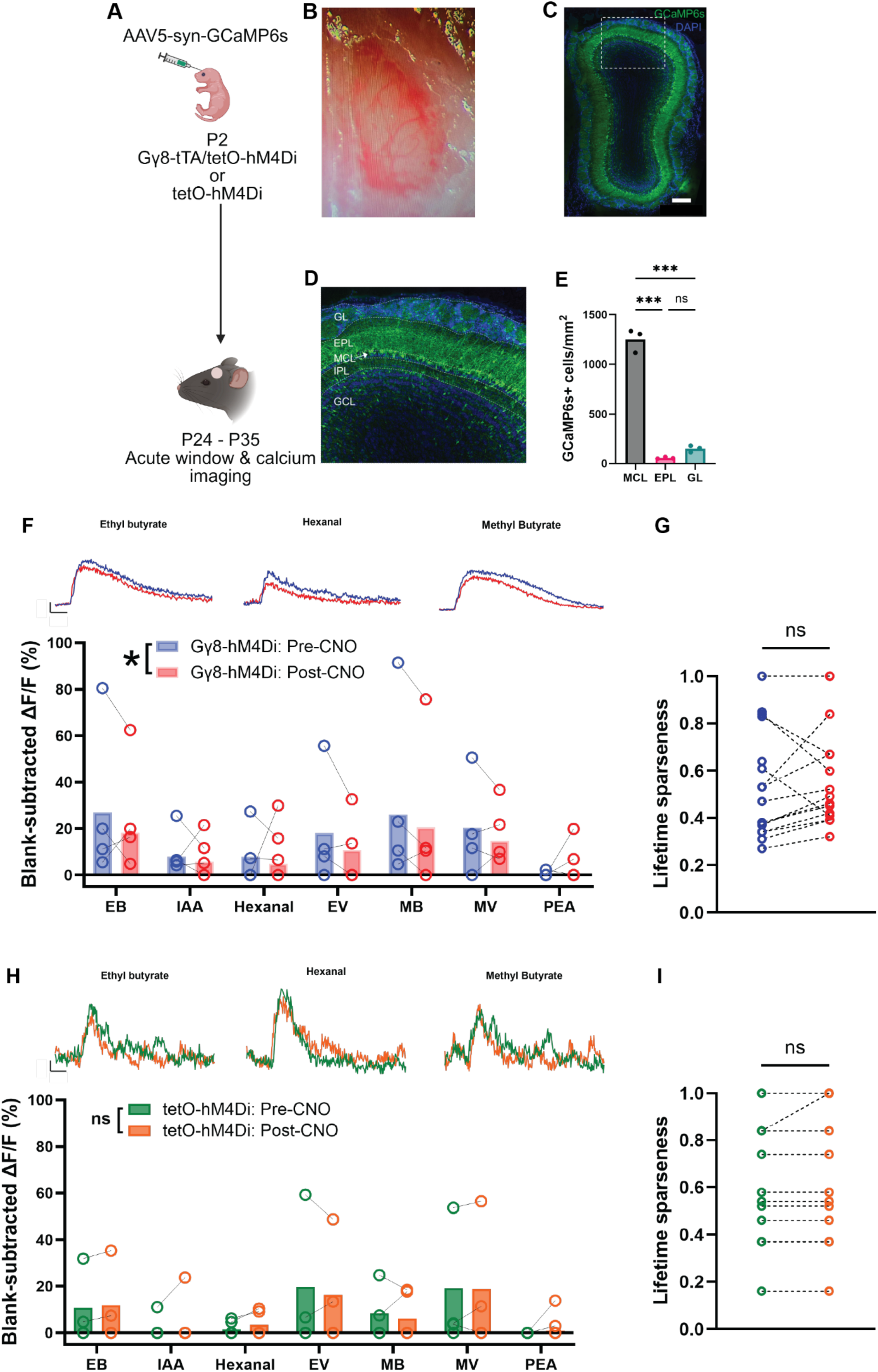
Acute silencing of immature OSN input dampens odor-evoked responses of postsynaptic cells across the OB. **A.** Neonatal injection strategy to introduce GCaMP6s into the OB for acute imaging. Gγ8-hM4Di (n = 4) and tetO-hM4Di (n = 4) mice received a 1 μL injection of AAV5-syn-GCaMP6s into the right OB at postnatal day 2 (P2). **B.** Acute cranial window placed over the right OB for 2-photon calcium imaging. **C.** Neonatal OB injection of AAV5-syn-GCaMP6s results in GCaMP6s expression. Stitched 10x magnification image of coronal section showing GCaMP6s expression throughout the entire OB. Scale bar = 250 µm. **D.** Inset of dorsal OB showing GCaMP6s expression is stronger in the MCL. IPL = internal plexiform layer, GCL = granule cell layer. **E.** Density of GCaMP6s+ cells in the MCL, EPL, and GL shows a majority of GCaMP6s expressing cells are found in the MCL, suggesting that odor-evoked calcium responses recorded in the GL are likely comprised of MC and TC dendritic responses (one-way ANOVA, *P* = 0.006, F2,6 = 13.7. Tukey’s multiple comparisons tests, MCL vs. EPL: q(6) = 7.34, *P* = 0.005. MCL vs. GL: q(6) = 2.88, *P* = 0.19. EPL vs. GL: q(6) = 4.46, *P* = 0.045. n = 3 mice). **F.** Acute silencing of immature OSNs via CNO injection decreases mean odor-evoked response amplitude in Gγ8-hM4Di mice (2-way ANOVA. Effect of CNO administration: *P* = 0.014, *F*(1,21) = 7.27. Effect of odors: *P* = 0.74, F(1,21) = 0.58. Interaction between odors and CNO administration: *P* = 0.78, F(1,21) = 0.53. *n* = 4 mice). Symbols: individual mice. Odors used in panel: ethyl butyrate [EB], isoamyl acetate [IAA], hexanal [HEX], ethyl valerate [EV], methyl butyrate [MB], methyl valerate [MV], and phenethylamine [PEA]. Traces above graphs (from left to right) are example traces of EB-, HEX-, and MB-evoked responses before and after CNO administration. Scale bar: 2 s (x), 25% (y). **G.** Lifetime sparseness of glomerular odorant responses is not affected by CNO administration in Gγ8-hM4Di mice (paired t-test. *P* = 0.44, *t*(13) = 0.80, n = 16 responding glomeruli from 4 mice). Symbols: values for individual glomeruli. **H.** CNO administration does not affect odor-evoked response amplitude in tetO-hM4Di control mice (2-way ANOVA. Effect of CNO: *P* = 0.096, F(1,21) = 3.04. *n* = 4 mice. Effect of odors: *P* = 0.80, F(6,21) = 0.50. Interaction between odors and CNO administration: *P* = 0.91, F(6,21) = 0.34. *n* = 4 mice). Odors, traces and symbols as in (**F**). **I.** CNO does not affect lifetime sparseness of odor-evoked responses in tetO-hM4Di mice (paired t-test. *P* = 0.34, *t*(10) = 1.00. *n* = 12 responding glomeruli from 4 mice). Symbols: values for individual glomeruli. Note that some symbols overlie each other in panels F and H, particularly for values of zero.

We found that on average, silencing immature OSNs significantly reduced the amplitude of odor-evoked glomerular responses across a panel of seven odorants (Fig. 3F, Videos S1, S2). Our findings in Fig. 3F are reported as mean odor-evoked response for each imaged mouse. When we analyzed the individual glomeruli that contribute to these per mouse mean responses, we saw a similar decrease in the amplitude of odor-evoked responses after CNO treatment (Fig. S2A). Hence, immature OSNs make a meaningful contribution to the odor information that is supplied to the OB in healthy adolescent mice. We also note that in both per glomerulus and per mouse data, occasional increases in odor-evoked response amplitude were observed after CNO treatment (Fig. 3F, Fig. S2A).

To further evaluate the selectivity of odor-evoked responses before and after immature OSN silencing, we calculated the lifetime sparseness (S_L_) of glomerular odorant responses. An S_L_ value of 1 indicates that a glomerulus responds to a single odorant (highly selective), whereas an S_L_ value of 0 indicates that a glomerulus responds equally to all seven odorants in our panel (non-selective). S_L_ values ranged from 0.27 to 1.0 for odor-responsive cells in Gγ8-hM4Di mice, indicating that some cells responded to multiple odors in our panel while others responded to only one (Fig. 3G). However, we found that odor selectivity was not impacted by immature OSN silencing, with no significant change in S_L_ after CNO treatment (Fig. 3G). CNO administration in tetO-hM4Di control mice did not impact the amplitude of odor-evoked responses *in vivo* (Fig. 3H, Fig. S1B). S_L_ values for cells in tetO-hM4Di mice were similar to those in Gγ8-hM4Di mice, and there was no significant change in S_L_ after CNO treatment in tetO-hM4Di mice (Fig. 3I). This suggests that CNO treatment does not affect the amplitude or odor selectivity of odor-evoked responses in mice that lack hM4Di expression. Overall, these results suggest that immature OSNs contribute to the odor input received by OB neurons in the healthy, intact olfactory system.

## Discussion

In this study we sought to define the contribution of immature OSNs to olfaction in the healthy, intact olfactory system. We showed that chemogenetically silencing immature OSNs in awake, behaving mice impairs odor detection ability. Furthermore, silencing immature OSNs results in decreased neuronal activity marked by immediate early gene expression in MCs throughout the OB and a broad reduction in the amplitude of odor-evoked responses across multiple glomeruli *in vivo* without affecting odor selectivity. Together, these data suggest that immature OSNs provide complementary odor input to that supplied by mature OSNs in the intact olfactory system.

An important question to address is what proportion of sensory input to the OB is being silenced by CNO treatment in Gγ8-hM4Di mice. While this value is difficult to determine definitively, it is possible to estimate an upper bound based on our findings in this study and data from previous studies. In our previous study^25^, we measured the linear density of mature OSNs expressing an OMP-driven reporter (36 per 100 μm) and immature OSNs expressing a Gγ8-driven reporter (20 per 100 μm) in 8-week-old mice). In this study, we found that Gγ8-driven hM4Di expression is present in only 60% of immature (GAP43-expressing) OSNs, meaning that GAP43+Gγ8- immature OSNs would account for an additional 13 cells per 100 μm. Therefore, Gγ8+ OSNs account for ∼29% of the total immature and mature OSN population. Note that this value excludes nascent OSNs, as they do not provide sensory input to the OB. In young adult mice, GAP43 expression begins as early as one day after terminal cell division, peaks at five days and is downregulated after eight days^14^. The onset of Gγ8 expression is unknown, but a conservative estimate would be two days after terminal cell division, based on the finding that Gγ8+ OSNs occupy slightly more apical positions in the OE than GAP43+ OSNs, indicating that they are slightly more mature^25^. Expression of Gγ8 is rapidly downregulated once OSNs reach eight days after terminal cell division^18^, suggesting a seven-day window (days 2-8 post-terminal cell division) for Gγ8 expression. The earliest time point at which there is anatomical and/or functional evidence for immature OSNs reaching the OB GL is at five days after terminal cell division^14,17,27^. Assuming that the rate of neurogenesis is relatively constant, only Gγ8+ OSNs that are 5-8 days post-terminal cell division could contribute to OB sensory input, i.e., 4/7 of the total Gγ8+ population. 4/7 of 29% = 17%, which is the maximum percentage of total OSN input that could be silenced by CNO treatment in Gγ8-hM4Di mice. Given this estimate, reducing immature OSN input to the OB seems to have surprisingly significant effects on odor detection behavior and odor-evoked responses, suggesting that their input may be distinct from that provided by mature OSNs.

We have shown previously that a subset of mice can perform odor detection and discrimination tasks, including the tasks used in the current study, using only immature OSNs following methimazole-mediated OSN ablation and subsequent regeneration of OSNs^27^. In that previous study, early recovery timepoints (5–7 days post-MMZ) at which mature OSNs are absent from the olfactory epithelium were tested^27^. However, an important remaining question was whether immature OSN activity is necessary for odor-guided behaviors in the healthy, intact olfactory system. Here, we demonstrated that silencing immature OSNs impairs odor detection ability in the buried food task (Fig. 2B), suggesting that immature OSNs may make a unique contribution to the odor information that is used to perform the buried food assay, which cannot be compensated for by mature OSNs. This raises the question of what this unique information may be, and our previous finding that immature OSNs encode odor concentration information high in the concentration range, when mature OSN responses have already saturated^27^, provides a potential explanation: immature OSNs may play an important role in odor source localization. Hence, sensory input from mature OSNs may play a more important role far from the buried food, whereas immature OSN input is important, or even necessary, to pinpoint the exact location.

Our data from the odor habituation-dishabituation task are difficult to interpret. The similar habituation and dishabituation indices in CNO- and saline-treated Gγ8-hM4Di mice (Fig. 2H,I), as well as the significant habituation in both CNO- and saline-treated Gγ8-hM4Di mice and significant dishabituation in CNO-treated Gγ8-hM4Di mice (Fig. S1A,C,D), suggest that immature OSN silencing may not affect odor discrimination in this assay. However, the lack of significant dishabituation in saline-treated Gγ8-hM4Di and of significant habituation or dishabituation in either tetO-hM4Di group raises questions about the utility of this assay as performed in this study. Therefore, we cannot draw strong conclusions from the odor habituation-dishabituation data. Possible sources of variability compared to our previous study^27^ include the younger age of mice^37^ (6–8 weeks at the time of testing) and the use of tea balls containing odorized filter paper rather than odorized cotton swabs. Additional studies will be needed to conclusively determine whether immature OSNs play a role in odor discrimination in the healthy, intact olfactory system, and could include a better optimized passive odor habituation-dishabituation assay and trained odor discrimination assays (e.g. go/no-go tasks of varying difficulty).

Our calcium imaging data showed that overall, silencing immature OSNs significantly reduces the amplitude of odor-evoked responses in postsynaptic OB neurons *in vivo*. However, there were some glomeruli for which odor-evoked response amplitude increased after CNO treatment. Our AAV5-based strategy to introduce GCaMP6s into OB neurons does result in expression throughout all layers of the OB, but GCaMP6s is preferentially expressed in the MCL and EPL of the OB (Fig. 3C–E). Thus, we reason that MC and TC dendritic responses are a major contributor to the GL GCaMP6s responses that we imaged. The effects of immature OSN silencing on GCaMP6s responses were similar across our odor panel, which includes the major odorant functional groups of esters (ethyl butyrate, isoamyl acetate, ethyl valerate, methyl valerate, and methyl butyrate), an aldehyde (hexanal), and an amine (phenethylamine) (Figs. 3F, S1A; Videos S1 & S2). Additionally, we sampled from a random population of glomeruli across the dorsal surface of the OB in each mouse. These factors together suggest that the changes in odor-evoked response amplitude elicited by immature OSN silencing are a generalized phenomenon in glomeruli populating at least the dorsal surface of the OB. This suggests that immature OSNs that project to the OB GL contribute to ongoing odor processing in the healthy olfactory system.

What mechanism(s) could underlie these changes in OB GL odor-evoked response amplitude following immature OSN silencing? Immature OSNs are known to form monosynaptic connections with external tufted cells^27^ and evoke firing in GL, EPL and MCL neurons *in vivo*^25^. Hence, a simple explanation for the overall reduction in odor-evoked response amplitude when OSNs are silenced is a reduction in excitatory afferent input to the imaged neurons. We also considered several possible non-mutually exclusive explanations for the less commonly observed increases in odor-evoked response amplitude post-CNO treatment. The first is that reduced afferent input also reduces lateral inhibition^38,39^ onto imaged MCs and TCs, with the net effect on an individual glomerulus depending on the odorant used and the pattern of excitatory and inhibitory inputs received by neurons that project their dendrites into that glomerulus. The second explanation involves relief of GABA_B_ and/or dopamine D2 receptor-mediated inhibition of OSN presynaptic terminals^40–43^ due to reduced excitatory input (whether mono-or di-synaptic) to dopamine- and GABA-releasing OB neurons. Finally, a modeling study suggested that ephaptic transmission between OSN axons could occur in tightly packed olfactory nerve fascicles^44^. This raises the theoretical possibility that immature OSN silencing could also reduce the activity of mature OSNs, which would result in greater changes in OB neuron activity. We note, however, that there is no experimental evidence for ephaptic transmission between mammalian OSN axons, nor could gap junctions between OSN axons, which would be required to mediate ephaptic transmission, be identified^45^.

We also found that immature OSN silencing did not affect the odor selectivity of odor-evoked responses in OB neurons (Fig. 3G). This finding agrees with our previous study, which found that the selectivity of odor-evoked responses was similar in immature and mature OSN axons^27^. In contrast, single cell RNA sequencing studies found that some immature OSNs express multiple OR transcripts^46,47^, which might be expected to reduce odor selectivity. The explanation for this discrepancy is provided by a recent study describing a Mex3a-dependent post-transcriptional regulatory mechanism that represses translation of multiple ORs in immature OSNs^48^.

A limitation of this study is that *in vivo* 2-photon calcium imaging was restricted to dorsal OB glomeruli, due to their optical accessibility. Hence, while silencing immature OSNs reduces the amplitude of dorsal GL odor-evoked responses, does silencing have a similar impact in other OB regions? Furthermore, mice were sedated using dexmedetomidine for imaging. While the half-life for dexmedetomidine elimination has not been measured in mice, it is ∼65 min in rats^49^, suggesting that sedation depth would be gradually decreasing during the ∼90 min time course of our imaging experiments. Given that odor representations are sparser in awake than in anesthetized mice^50,51^, changes in sedation depth could have affected the amplitude of odor-evoked responses over time.

Another limitation was that we did not obtain direct evidence of CNO-mediated silencing from *in vivo* recording of OSNs in Gγ8-hM4Di mice. Nevertheless, we did obtain functional evidence for the efficacy of immature OSN silencing from immediate early gene staining of odor-evoked MC activity, which shows a significant reduction in the number of PS6+ cells throughout the dorsal and ventral halves of the OB (Fig. 1G-I). This was surprising in light of previous studies showing that ventral OE regions have more proliferating cells and a greater proportion of immature OSNs than dorsal OE regions^17,34^, which might be expected to result in greater immature OSN innervation of ventral vs. dorsal OB. While more sensitive assays of neuronal activity, that also assess other OB neuron types, might detect such a difference, the key finding here is that immature OSN input is sufficient throughout the OB to impact MC activity when it is silenced.

Together, our data highlight the important contribution of postnatally born OSNs to normal odor processing even while these cells are in development. It was once believed that newborn OSNs did not functionally contribute to olfactory circuity until they had reached maturity. In contrast, we have shown that immature OSNs not only contribute to odor-evoked responses but also provide odor input that is useful in detecting odors in behaving mice with a healthy, intact olfactory system and cannot be supplied by mature OSNs. Hence, we reason that immature OSNs provide complementary input to that supplied by mature OSNs that is essential for olfactory processing and odor-guided behaviors that are important for survival.

## Methods

### Experimental animals

All animal procedures conformed to the National Institutes of Health guidelines and were approved by the University of Pittsburgh Institutional Animal Care and Use Committee. Mice were housed on a 12-hour light/dark cycle in individually ventilated cages at 22°C and 48% humidity with unrestricted access to food and water unless otherwise stated. Mice were group-housed if same-sex littermates were available. A total of 59 mice was used in this study.

Generation of the Gγ8-tTA^52^ and tetO-hM4Di^31^ (Jackson Laboratory strain #024114, https://www.jax.org/strain/024114) mouse lines has been published. Separate cohorts of experimental mice were used for the following: behavioral testing (buried food and habituation/dishabituation tests) (age range: 6 to 8 weeks), *in vivo* 2-photon calcium imaging (age range: postnatal days 24 to 35 [P24-35]) and immunohistochemistry (age range: 6 to 8 weeks). All mice used were either Gγ8-hM4Di [Gγ8-tTA^+/-^;tetO-hM4Di^+/-^] or tetO-hM4Di^+/-^ genotypes. Each experimental group comprised roughly equal numbers of male and female mice. Mice of each genotype were randomly assigned to each experimental group for behavioral testing. Mice were genotyped by PCR using previously validated primers as described previously^27^

(https://www.jax.org/Protocol?stockNumber=024114&protocolID=26393).

### Clozapine-N-Oxide (CNO) treatment to silence immature OSNs

30 minutes prior to starting behavioral or 2-photon calcium imaging experiments (to allow for CNO to begin taking effect), mice received a single intraperitoneal injection of either saline (vehicle) or CNO (1 mg/kg; 10 μl/g). All experiments were conducted 30 minutes after injection to allow for the CNO to have taken effect well, but remain within its ∼1-hour half-life^53,54^.

### Odor exposure assay for immediate early gene phospho-S6 immunohistochemistry

30 minutes prior to odor exposure, mice (age range: 6 to 8 weeks) received an i.p. injection of either CNO or saline and were housed in a separate empty home cage to allow the drug to take effect. After these 30 minutes passed, mice were then placed individually into new empty home cages with the cage lid on and were exposed for 1 hour to a glass vial with a perforated lid containing a mixture composed of 30 μL each of the following odorants, diluted 1:10 in mineral oil: isoamyl acetate, ethyl butyrate, methyl valerate, and isovaleric acid. Mice underwent transcardial perfusion immediately after the 1-hour exposure.

### Transcardial perfusion and cryosectioning

Mice were deeply anesthetized using 4% isoflurane in 1L/min O_2_ and transcardially perfused with phosphate buffered saline (PBS) followed by 4% paraformaldehyde (PFA). The OBs of these mice were then dissected out, cryoprotected in a 30% sucrose solution, embedded in 10% gelatin, fixed/cryopreserved, flash frozen, and sectioned on a cryostat for immunostaining as described previously^27^.

### Immunohistochemistry in OB and OE sections

Free floating OB and OE sections were treated with 1% sodium borohydride, washed with PBS, and then permeabilized for one hour in blocking buffer (5% normal donkey serum [NDS]/0.5% Triton X-100 in PBS). 40 μm OB sections were then stained with a primary antibody to detect the immediate early gene (IEG) phospho-S6 (PS6). 40 μm OB sections from mice that had been neonatally injected with AAV5-syn-GCaMP6s were stained with anti-GFP-ATTO488 nanobody. 50 μm OE sections were co-stained with primary antibodies to detect immature OSNs via growth-associated protein 43 (GAP43)^11^ and the hM4Di receptor via its hemagglutinin (HA) tag. All sections were incubated in primary antibody solutions (3% NDS, 0.2% Triton X-100, 0.01% sodium azide in PBS) as shown in Table 1. Following primary antibody application, sections were washed in PBS and then incubated in secondary antibody solution (3% NDS, 0.2% Triton X-100 in PBS) for one hour at room temperature (except for ATTO-488-GFP staining, which did not require secondary antibody incubation). Sections were then washed in PBS and mounted with Vectashield containing DAPI (Vector Labs).

**Table 1:** Primary and secondary antibody information.

| Antibody | Dilution | Species | Incubation conditions | RRID | Manufacturer catalog # |
| --- | --- | --- | --- | --- | --- |
| GAP43 | 1:500 | Mouse | 96 h \| 4 °C | AB_2533123 | ThermoFisher 33-5000 |
| Anti-HA | 1:250 | Rabbit | 96 h \| 4 °C | AB_1549585 | Cell Signaling Technology #3724 |
| Phospho-S6 | 1:400 | Rabbit | 96 h \| 4 °C | AB_916156 | Cell Signaling Technology #4858 |
| ATTO-488-GFP | 1:500 | Camelid | 1 h \| room temp. | AB_2631386 | Chromotek #gba488 |
| Rabbit-AF488 | 1:500 | Donkey | 1 h \| room temp. | AB_2535792 | ThermoFisher A21206 |
| Rabbit-AF546 | 1:500 | Donkey | 1 h \| room temp. | AB_2534016 | ThermoFisher A10040 |
| Mouse-AF546 | 1:500 | Donkey | 1 h \| room temp. | AB_11180613 | ThermoFisher A10036 |

### Confocal microscopy

Images of stained OE sections were acquired using an Olympus Fluoview 3000 confocal microscope with an Apo 60x/1.4 NA oil-immersion objective and Fluoview imaging software (Olympus). Laser excitation and bandpass emission filters were: 405 nm and 450/50 for DAPI, 488 nm and 525/50 for AF488, and 561 nm and 595/50 for AF546. Z series images were collected with 2 μm steps between each z plane. Images were 212.1 x 212.1 μm (1024 x 1024 pixels) collected with a 2.0 μs/pixel sampling speed.

### Widefield fluorescence microscopy

Images of OB sections were collected using Echo imaging software (Discover Echo) on a Revolve microscope equipped with LED excitation (Discover Echo) and a Plan Apo 10x/0.4 NA air objective (Olympus). Images were stitched together in Affinity Photo software (Affinity). Bandpass emission filter wavelengths were (in nm): DAPI (450/50), GFP (495/50) and Phospho-S6-AF546 (590/50). Detection settings were identical for all images collected for this dataset.

### Image analysis in OB and OE sections

Image analysis for PS6+ cell counts in OB sections, colocalization of GAP43 and HA in OE sections and GCaMP6s expression in OB sections was performed in Fiji (ImageJ)^55^. Images of OB sections were subdivided into dorsal and ventral halves by horizontally bisecting the image halfway along a vertical line extending from the center of the dorsal surface to the center of the ventral surface. MCs, defined as large neurons within the MCL, that expressed PS6 were counted and the length of the MCL/EPL border was measured, for dorsal and ventral halves of each section. PS6+ MC linear density (MCs per mm) was then calculated separately for each half, and as a total value for the entire section. Cells expressing GCaMP6s (as marked by anti-GFP staining) were counted in the MCL, EPL and GL, and their density in each layer was calculated. OSNs expressing either GAP43 or HA were marked separately in z-stacks. The proportion of immature OSNs in the OE that expressed the hM4Di receptor was then calculated by dividing the number of co-stained (GAP43+ and HA+) cells by the total number of immature (GAP43+) OSNs and converting it to a percentage.

### Odor detection and discrimination assays and analysis

A buried food assay^27,56^ was used to test odor detection ability following either CNO or saline injection, using a separate group of mice for these two experimental conditions. Two days before testing, a Froot Loop (Kellogg’s) was placed into the home cage to familiarize the mice with the target odor. 14 hours prior to testing, mice were placed in a new home cage and food restricted until testing. During testing, 6–8-week-old mice were placed in an empty cage (L: 29.21 cm x W: 19.05 cm x H: 12.7 cm) with 1.5 cm of autoclaved bedding for five minutes to acclimate to the environment. Mice were then removed so that the Froot Loop could be buried in the center of the cage. The mice were then placed back into the cage and given 10 minutes to complete the task. Their behavior during both the acclimation and testing periods was monitored via video recording software (pylon Viewer, Basler). The amount of time spent digging during the five-minute acclimation period was recorded for each mouse. The time that it took for the mouse to uncover the buried food was reported. Mice failed the task if they did not uncover the buried food within 10 minutes. If mice failed to complete the task within the allotted ten minutes, they were removed from the cage, assigned a time of 10 minutes, and given the cereal to eat in their home cage. Mice that uncovered the buried food within the allotted 10 minutes of the test were allowed to eat the cereal before being returned to their home cage.

The day after the buried food assay, an odor habituation-dishabituation assay was used to test odor discrimination ability in the same saline or CNO-injected mice, using some modifications from previous protocols^27,56^. Mice were acclimated for 30 minutes to the test cage, which was an empty cage (length: 30.48 cm x width: 19.05 cm x height: 12.7 cm) with a metal tea ball secured to the center of the cage floor. Mice were then presented with a series of odorant-soaked filter paper squares placed in the tea ball for two minutes each. The series of odorants consisted of three 2-minute trials each of water, almond extract, mint extract, and vanilla extract (all diluted 1:100 in water). Live recordings were captured via pylon Viewer software, and investigation of the tea ball during each odor presentation was analyzed using EthoVision tracking software (Noldus). Time spent investigating the tea ball for each odor and total investigation time across the entire test period were calculated. A habituation index value was calculated by subtracting the amount of time the mouse spent sniffing during the second trial from the first trial of each odorant. A dishabituation index value was calculated by subtracting the amount of time the mouse spent sniffing the third presentation of an odorant from the amount of time they spent sniffing the first presentation of the subsequent odorant.

### Neonatal AAV injections in mouse olfactory bulbs

At P2, Gγ8-hM4Di and tetO-hM4Di mice were anesthetized by acute hypothermia^57^ until immobile and 1 μL of AAV5-syn-GCaMP6s (Addgene #100843-AAV5) was injected into the right OB via a glass micropipette^58^. Mice then recovered on a heating pad until awake and mobile in a separate holding container before being returned to their home cage until they reached imaging age (P24-P35).

### Acute window preparation for *in vivo* 2-photon calcium imaging

P24-P35 Gγ8-hM4Di mice were anesthetized with isoflurane (1.5–2% in 1L/min oxygen), administered ketoprofen (5 mg/kg) as an analgesic, and received a ∼1 mm diameter craniotomy over the right OB where a glass coverslip cranial window was implanted^59^ (Fig. 3A & B). A head bar was also attached to the skull between the coronal and lambdoid sutures.

### *In vivo* 2-photon calcium imaging and analysis

After acute cranial window implantation and head bar affixation, mice received dexmedetomidine (0.5 mg/kg; 10 μL/g) for sedation while undergoing imaging. Mice were imaged using ThorImage 4.1 software (ThorLabs) and a Bergamo II 2-photon microscope with a resonant scanner and GaAsP detectors (ThorLabs) using a Semi/Apo 20x/1.0NA water-immersion objective (Olympus), and an Insight X3 IR laser (Newport) mode-locked at 935 nm to image GCaMP6s fluorescence. An imaging field of view was selected through GCaMP6s expression identified via widefield 490 nm LED (ThorLabs) illumination across the region of the dorsal OB surface visible through the cranial window.

Odorant stimulation was performed using a custom-built Arduino-controlled olfactometer. Stimulation and image acquisition timing were controlled and recorded using Python, ThorSync, and ThorImage (ThorLabs). A panel of seven odorants known to stimulate dorsal surface OB glomeruli was used. The following odorants were used at a 10% concentration diluted in mineral oil (v/v): ethyl butyrate (EB), isoamyl acetate (IAA), hexanal, ethyl valerate (EV), methyl butyrate (MB), methyl valerate (MV), and 2-phenethylamine (PEA). Saturated vapor from odorant vials, or a blank deodorized air control, were delivered via an oxygen carrier stream (1 L/min flow rate). All stimuli were 1 s in duration and were delivered via a tube positioned ∼5 mm from the mouse’s nose. All image trials were 20 s in duration (3.5 s baseline, 1 s stimulus, 15.5 s post-stimulus). Three trials were collected for each stimulus, with odorants and blank stimuli delivered in a pseudo-randomized order with a 60 s inter-trial interval.

Images were analyzed in Fiji. Glomeruli were manually outlined in each field of view, and these outlines were used for analysis of all stimulus-evoked responses. Fluorescence intensity over time was calculated for each glomerulus for each trial. Mean values from the three trials for each stimulus were then calculated for each glomerulus. Additionally, the mean value across all glomeruli was calculated for each mouse. The fluorescence intensity change (ΔF) was calculated as the difference between a 1 s baseline prior to stimulus onset and peak fluorescence within 3 s after stimulus onset. A glomerulus was defined as responding to a stimulus if the Δ*F* amplitude was greater than three standard deviations of the baseline period. ΔF/F (%) was reported as (peak - baseline)/baseline x 100 for significant responses only. For blank subtraction, the ΔF/F in response to the blank (deodorized air) stimulus for a glomerulus was subtracted from the ΔF/F in response to each odorant.

Lifetime sparseness (S_L_) was calculated to measure the degree of odor selectivity of each glomerulus before and after CNO treatment. S_L_ was calculated as (1 − (Σ_i=1,n_(*r_i_*/*n*)^2^)/(Σ_i=1,n_(*r_i_*^2^/*n*)))/(1 − 1/*n*), where *r_i_* is the ΔF/F of the glomerular response to odorant *i* and *n* is the total number of odorants (Huang et al., 2022). A highly odorant-selective glomerulus had an S_L_ of 1 and a glomerulus that was equally responsive to all odors in the panel had an S_L_ of 0.

### Statistics

Statistical analyses were conducted in Prism 9 or Prism 10 software (GraphPad). All data were tested for normality (Shapiro-Wilk or Kolmogorov-Smirnov test) and equal variance (Browne-Forsyth test) to determine whether parametric tests could be employed. Co-staining for the HA tag on the hM4Di receptor in immature OSNs (identified via GAP43 expression) in olfactory epithelia of Gγ8-hM4Di and tetO-hM4Di mice was assessed using an unpaired t-test. Linear density of PS6+ cells in saline- vs. CNO-treated Gγ8-hM4Di and tetO-hM4Di mice was compared using unpaired t-tests. For the buried food task, Welch’s one-way ANOVA with Dunnett’s T3 multiple comparisons tests were conducted on the time to success and on digging time during the acclimation period. For the habituation/dishabituation task, Welch’s one-way ANOVA with Dunnett’s T3 multiple comparisons tests were used to compare total investigation time, habituation index and dishabituation index between saline- and CNO-treated Gγ8-hM4Di and tetO-hM4Di mice. We also determined whether each group showed significant habituation and/or dishabituation by comparing habituation and dishabituation indices to zero using paired t-tests. The density of GCaMP6s+ cells across OB layers was compared using a one-way ANOVA with Tukey’s multiple comparisons tests. Odor-evoked calcium responses before and after CNO treatment were analyzed using two-way ANOVAs. Finally, paired t-tests were used to compare lifetime sparseness before and after CNO treatment.

## Acknowledgements

Research reported in this manuscript was supported by the National Institute on Deafness and other Communication Disorders of the National Institutes of Health under Award Number R01DC018516 to C.E.J.C. The content is solely the responsibility of the authors and does not necessarily represent the official views of the National Institutes of Health. We thank members of the Cheetham lab for helpful discussions.

## Data availability

The data from this manuscript are available at: https://doi.org/10.5281/zenodo.19235831

## Supplemental Material

### Supplemental Figures

**Figure S1.**
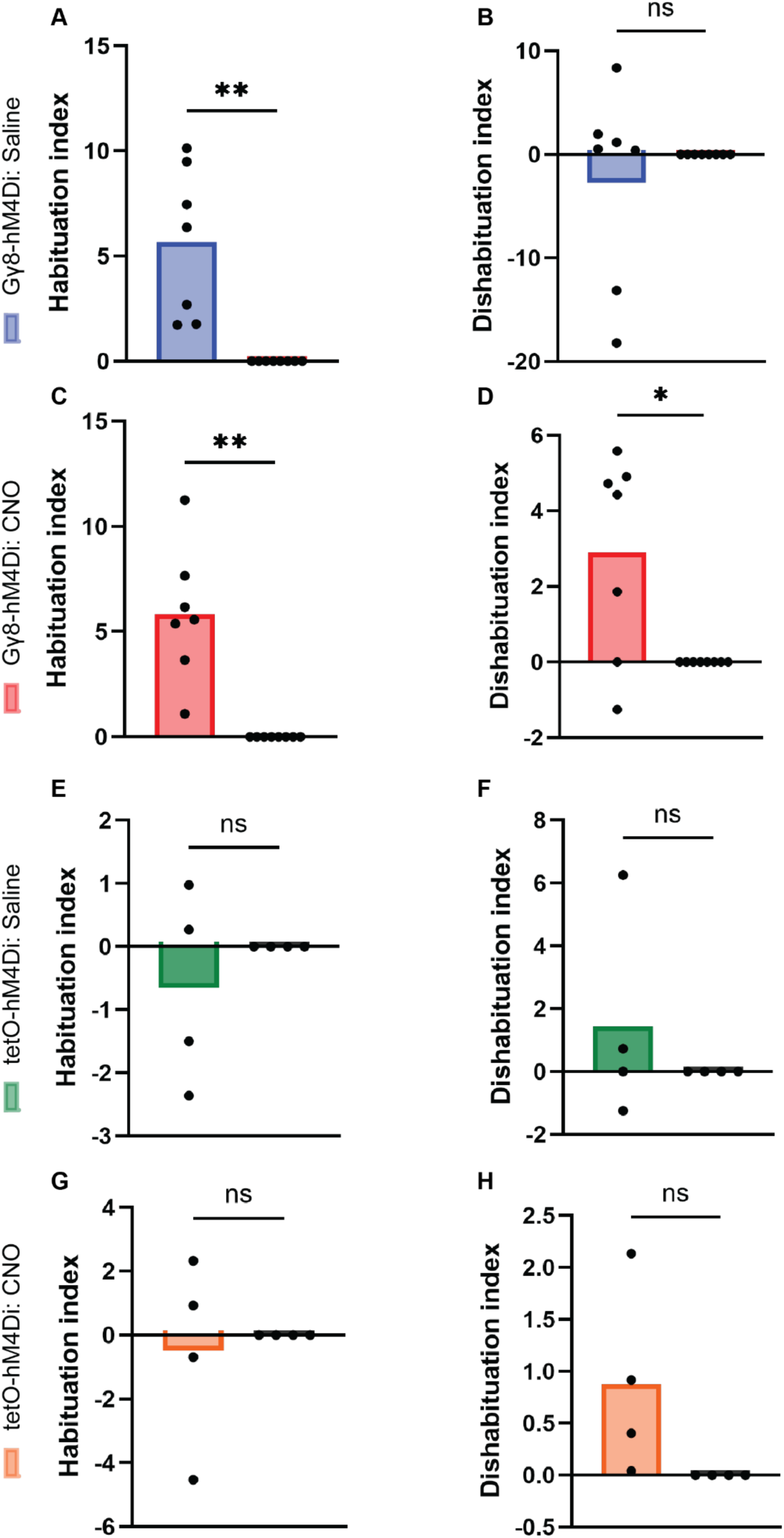
Testing experimental group habituation and dishabituation indices against zero reveals variable performance in the odor habituation-dishabituation assay. Habituation or dishabituation index values for each experimental group were compared to zero using Welch’s paired t-tests to determine whether mice in that group showed significant habituation or dishabituation. **A.** Habituation index in saline-treated Gγ8-hM4Di mice (t (6.00) = 4.16, *P* = 0.006). **B.** Dishabituation index in saline-treated Gγ8-hM4Di mice (t (6.00) = 0.77, *P* = 0.47). **C.** Habituation index in CNO-treated Gγ8-hM4Di mice (t (6.00) = 4.86, *P* = 0.003). **D.** Dishabituation index in CNO-treated Gγ8-hM4Di mice (t (6.00) = 2.84, *P* = 0.030). **E.** Habituation index in saline-treated tetO-hM4Di mice (t (3.00) = 0.85, *P* = 0.46). **F.** Dishabituation index in saline-treated tetO-hM4Di mice (t (3.00) = 0.87, *P* = 0.45). **G.** Habituation index in CNO-treated tetO-hM4Di mice (t (3.00) = 0.33, *P* = 0.76). **H.** Dishabituation index in CNO-treated tetO-hM4Di mice (t (3.00) = 1.91, *P* = 0.152). Symbols: individual mice or zero values for analysis.

**Figure S2.**
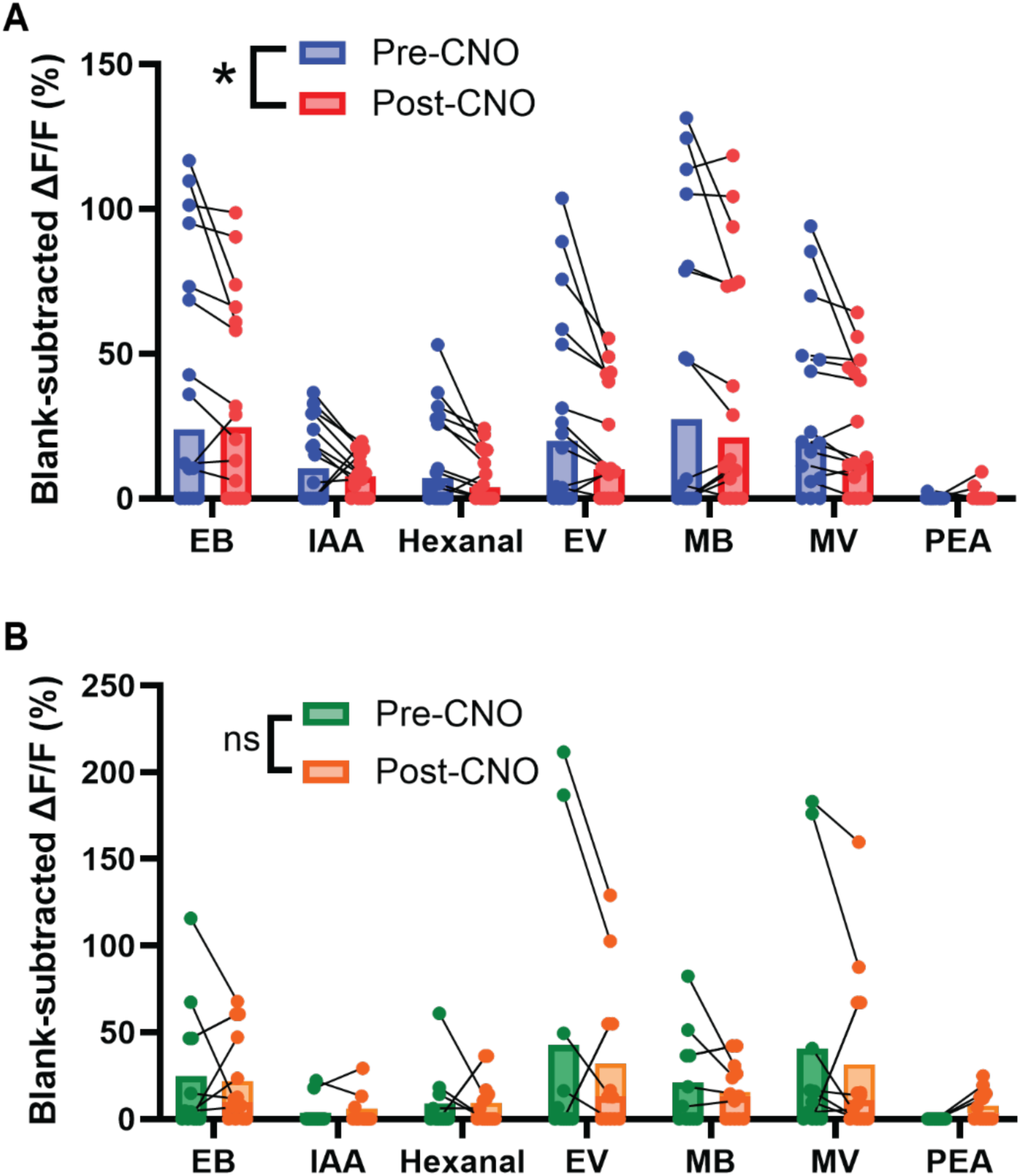
Acute silencing of immature OSN input dampens odor-evoked responses of postsynaptic cells on a per-glomerulus basis. **A**. Acute silencing of immature OSNs via CNO injection reduces mean odor-evoked responses in the individual glomeruli of Gγ8-hM4Di mice generally (2-way ANOVA. Effect of CNO administration: *P* < 0.001, F_1,105_ = 22.8. Effect on odors: *P* < 0.001, F_6,105_ = 5.06. Interaction between odors and CNO administration: *P* = 0.34, F_6,105_ = 1.15, n = 16 responsive glomeruli from 4 mice). **B**. CNO administration has no effect on odor-evoked responses in the glomeruli of tetO-hM4Di control mice generally (2-way ANOVA. Effect of CNO administration: *P* = 0.43, F_1,68_ = 0.62. Effect on odors: *P* =0.081 F_6,86_ = 1.96. Interaction between odors and CNO administration: *P* = 0.90, F_6,68_ = 0.37, n = 14 responsive glomeruli from 4 mice). Odors used in panel: ethyl butyrate [EB], isoamyl acetate [IAA], hexanal [HEX], ethyl valerate [EV], methyl butyrate [MB], methyl valerate [MV], and phenethylamine [PEA].

### Supplemental Video Legends

**Supplemental Video 1. Glomerular layer odor-evoked responses in OB neuron dendrites.** Representative video showing ethyl butyrate-evoked responses captured in the glomerular layer of the OB during odor presentation before CNO administration. Here, we see the emergence of ethyl butyrate responses in both dendritic processes throughout the glomeruli and surrounding cell bodies. Responses are shown in real time (15 frames per s).

**Supplemental Video 2. Silencing immature OSNs reduces the amplitude of glomerular layer odor-evoked responses in OB neuron dendrites within glomeruli.** Representative video of ethyl butyrate-evoked responses in the same field of view as in Supplemental Video 1 after CNO administration to silence immature OSNs. Odor-evoked responses in dendrites within the imaged glomeruli are reduced in amplitude relative to before CNO treatment.

